# Theaflavins, polyphenols of black tea, inhibit entry of hepatitis C virus

**DOI:** 10.1101/325126

**Authors:** Pritom Chowdhury, Marie-Emmanuelle Sahuc, Yves Rouillé, Alexandre Vandeputte, Priscille Brodin, Manoranjan Goswami, Tanoy Bandyopadhyay, Jean Dubuisson, Karin Séron

**Affiliations:** Univ. Lille, CNRS, INSERM, CHU Lille, Institut Pasteur de Lille, U1019 - UMR 8204 - CIIL - Center for Infection and Immunity of Lille, Lille, France; Department of Biotechnology, Tocklai Tea Research Institute, TRA, Jorhat, Assam, India; Department of Biochemistry, Tocklai Tea Research Institute, TRA, Jorhat, Assam, India

## Abstract

The treatment of hepatitis C virus (HCV) infection by combination of direct acting antivirals (DAA), with different mode of action, has made substantial progress in the past few years. However, appearance of resistance and high cost of the therapy is still an obstacle in the achievement of the therapy, more specifically in developing countries. In this context, search for affordable antivirals with new mechanisms of action is still needed. Tea, after water, is the most popular drink worldwide. Polyphenols extracted from green tea have already shown anti-HCV activity as entry inhibitors. Here, three different theaflavins, theaflavin (TF1), theaflavin-3’-monogallate (TF2), and theaflavin-3-3’-digallate (TF3), which are major polyphenols from black tea, were tested against HCV in cell culture. The results showed that all theaflavins inhibit HCV infection in a dose-dependent manner in an early step of infection. Results obtained with HCV pseudotyped virions confirmed their activity on HCV entry and demonstrated their pan-genotypic action. No effect on HCV replication was observed by using HCV replicon. Investigation on the mechanism of action of black tea theaflavins showed that they act directly on the virus particle and are able to inhibit cell-to-cell spread. Combination study with inhibitors most widely used in anti-HCV treatment regimen demonstrated that TF3 exerts additive effect. In conclusion, theaflavins, that are present in high quantity in black tea, are new inhibitors of HCV entry and hold promise for developing in therapeutic arsenal for HCV infection.

## Introduction

Hepatitis C caused by hepatitis C virus (HCV) has been called silent epidemic. The majority of infections are asymptomatic, but in 20% of cases the virus persist, leading to chronic hepatitis (1) causing liver fibrosis and cirrhosis, which is often a prelude to hepatocellular carcinoma (2). Liver transplantation is often necessary in a portion of HCV infected patients (3). Tremendous efforts have been expended to develop efficacious prophylactic and therapeutic treatment regimen for chronic hepatitis C. No vaccine is available due, at least in part, to the high genomic variability of HCV, which has led to the distinction of seven genotypes, most of which have multiple subtypes (4). The therapeutic option against HCV has recently been improved with the development of HCV direct acting antivirals (DAA) like Daclatasvir, Sofosbuvir and Simeprevir, targeting viral proteins NS5A, NS5B polymerase or NS3/4A protease, respectively (5). These approved DAA prominently increase the sustained viral response (SVR) up to ~95% in most patients, depending primarily on disease stage and the genotype of the infecting virus (5). However, treatment with DAAs is not without limitation; it is associated with side-effects, resurgence of infection in transplant patient and high cost especially in developing countries (6,7). Approved DAAs mainly target the virus replication leading to emergence of resistance mutations in this RNA virus genome (8). Thus, novel combinations of low cost entry inhibitors with conventional treatment targeting different stages of the HCV life cycle, may provide a promising approach against HCV drug resistance development and infection relapse (9). Moreover, prevention of donor liver re-infection by inhibiting viral entry into hepatocytes might be achieved using DAAs targeting entry.

Hepatitis C virus is an enveloped positive-stranded RNA virus encoding a polyprotein, co- and post-translationnally cleaved into structural and non-structural proteins (10). Two viral glycoproteins E1 and E2 are part of the lipoviroparticle envelope. Non-structural proteins, NS2 to NS5B, are involved in replication and assembly of new virions. Actual antiviral therapy with DAA targets three non-structural proteins, the RNA-dependent RNA polymerase NS5B, a non-enzymatic protein involved in replication and assembly of HCV NS5A, and the viral protease NS3/4A, involved in polyprotein processing and essential for viral replication (11). Virus entry into hepatocytes is a multistep process that involves attachment of the particle to glycosaminoglycans and subsequent binding to entry factors, SR-B1, CD81, Claudin-1 and Occludin (12). After clathrin-mediated endocytosis and fusion of the viral envelope to endosomal membrane, the viral RNA is replicated, assembled and released via the secretory pathway.

In recent times, a wide variety of natural compounds have been extensively studied in terms of their antiviral activity (13). Polyphenols are one such group of compounds with potent antiviral activities. Earlier studies of others and our group have shown that epigallocatechin-*3*-gallate (EGCG), a major green tea polyphenol, inhibits HCV entry by a new mechanism of action (14–16). Recently, black tea polyphenols, theaflavins (TFs), components of black tea extract, have shown potent antiviral activities against calcivirus (17), herpes simplex virus 1 (HSV-1) (18), human immunodeficiency virus 1 (HIV-1) (19) and influenza A (20). Theaflavins are formed by the enzymatic oxidation and decarboxylation of catechins during the manufacture of black tea (21). The most abundant TFs are theaflavin (TF1), theaflavin-3’-monogallate (TF2) and theaflavin-3-3’-digallate (TF3). Theaflavins share many of the structural characteristics of EGCG, which prompted us to screen theaflavins against HCV. We also aim to identify compounds with enhanced anti-HCV activity than previously studied compounds to boost the development of suitable inhibitors.

Here, we report the capacity of theaflavins to inhibit HCV infection *in vitro*, demonstrating their activity on HCV entry. Given the abundance of theaflavins, especially in tea growing areas, and popularity of tea as a drink next only to water and as health drink, the compounds hold promise for developing in therapeutic arsenal for HCV infection.

## Materials and Methods

### Chemicals and antibodies

DMEM, goat and fetal bovine sera were purchased from Invitrogen (Carlsbad, CA). 4’,6-diamidino-2-phenylindole (DAPI) was from Molecular Probes (Invitrogen). EGCG used as a control was from Calbiochem (Merck Chemicals, Darmstadt, Germany). Theaflavin 3,3’-digallate used to compare with our extracted theaflavin, TF3, was from PhytoLab (Germany). Boceprevir was kindly provided by Philippe Halfon (Hôpital Européen, Laboratoire Alphabio, Marseille, France). Sofosbuvir (PSI-7977) and Daclatasvir (BMS-790052) were purchased from Selleckchem (Houston, USA). All other chemicals like DMSO were purchased from Sigma. Mouse anti-E1 monoclonal antibody (MAb) A4 was produced *in vitro* (22). Cy3-conjugated goat anti-mouse IgG was from Jackson Immunoresearch (West Grove, PA).

### Cells and virus strains

Human hepatoma Huh-7 cells and human embryonic kidney HEK 293T cells were grown in Dulbecco’s modified Eagle’s medium (DMEM) supplemented with glutamax-I and 10% fetal bovine serum.

Japanese fulminant hepatitis-1 (JFH-1) HCV strain containing titer enhancing mutations (23) was used. Virus stocks were prepared by infecting Huh-7 cells with HCV JFH-1 (HCVcc) with a titer of 10^6^ pfu/ml.

### Theaflavins extraction and stock preparation

Crude theaflavin was extracted from prepared “crush tear curl” (CTC), black tea from Assam, India. The crude theaflavin in ethyl acetate was subjected to sephadex LH-20 column (Sigma) and eluted with 40% acetone solution. The elute with distinguished reddish colours for each of TF1, TF2 and TF3 was fractionated. Each fraction was concentrated by a rotary evaporator and lyophilized, and the three fractions of theaflavins recrystallized from dehydrated ethanol. The purity of the compounds was confirmed by HPLC, which was found to be up to 98%. Purified TF1, TF2, and TF3 were each dissolved in DMSO to produce stock solutions at a concentration of 250 mg/ml.

### Cytotoxicity assay

Huh-7 cells were seeded in 96 well plates and incubated at 37°C for 24 h. Cultures of 60-70% confluency were treated with various concentrations of theaflavins. An MTS [3-(4, 5-dimethylthiazol-2-yl)-5-(3-carboxymethoxyphenyl)-2-(4-sulfophenyl)-2H-tetrazolium, inner salt] based assay (CellTiter 96 aqueous nonradioactive cell proliferation assay, Promega) was used to evaluate cell viability treated after 24, 48 and 72 h. DMSO was used as control.

### Immunofluorescence assay for anti-HCV quantification in confocal microscopy

Huh-7 cells were seeded in 96-well plates (6.000 cells/well) the day before infection. Cells were infected for 2 h with HCVcc at a multiplicity of infection (MOI) of 0.8. Then, the inoculum was removed and cells were overlaid with fresh medium. After 30 h, cells were fixed with ice-cold methanol and were processed for immunofluorescent detection of E1 envelope protein as described (24). Nuclei were stained with 1 μg/ml DAPI. Confocal images were recorded using an automated confocal microscope InCell Analyzer 6000 (GE Healthcare Life Sciences). Each image was then processed using the Colombus image analysis software (Perkin Elmer) as described (14).

### Time of addition assay

Compounds were added before, during and after inoculation of Huh-7 cells with HCVcc. Briefly, cells were incubated in the presence or absence of compounds (pretreatment) for 1 h and subsequently replaced with virus for 1 h (inoculation). Inoculum was removed and replaced with DMEM for 28 h (post-inoculation). After 30 h cells were fixed and immunofluorescence detection was performed as described above.

### Direct effect on viral particle

HCV was pre-incubated with 25 μg/ml of theaflavins for 1 h before inoculation, and then the mixture was diluted 10 times before being used for the inoculation (1 h at 37°C), leading to a final concentration of theaflavins of 2.5 μg/ml. In parallel, Huh-7 cells were inoculated with HCV in the presence of theaflavins either at 2.5 or 25 μg/ml. Importantly, the MOI was kept constant in all the conditions. Inoculum was removed and replaced with DMEM. After 30 h, immunofluorescence was performed as described above.

### Entry assay with HCV pseudo-particles (HCVpp)

The luciferase-based HCV pseudotyped retroviral particles were generated as previouslydescribed (25). Briefly, HEK-293T cells were transfected with plasmids encoding HCV envelope proteins, Gag/Pol, and firefly luciferase. The HCV envelope plasmids included 6 different genotypes (1b, 2a, 3a, 4, 5 and 6). The supernatants of the transfected cells were collected 72 h later and filtered through a 0.45-μm membrane. Vesicular stomatitis virus pseudoparticles (VSVpp) were used as a control. For the entry assay, Huh-7 cells were seeded on 96-well plates overnight and incubated with HCVpp for 2 h at 37°C in the presence or not of theaflavins. Inoculum was replaced by DMEM and kept for 48 h. HCVpp entry into Huh-7 cells was measured by luciferase activity quantification using Luciferase Assay Kit (Promega) and a Berthold luminometer.

### HCV replicon and replication assay

The plasmid pSGR-JFH1 encoding a sub-genomic replicon of JFH-1 strain was obtained from Dr T Wakita (26). A *Bgl*II and an *Nsi*I restriction sites were inserted between codons Pro419 and Leu420 of NS5A, and the coding sequence of enhanced green fluorescent protein (EGFP) was then inserted between these two sites. This position was previously shown to accept a GFP insertion in a sub-genomic replicon of the Con1 strain (27). The plasmid was in vitro transcribed before electroporation into Huh-7 cells. Cells that express replicon were selected for using 500 μg/mL of geneticin during 15 days and cultured in a medium containing 250 μg/mL of geneticin. Huh-7 cells stably expressing replicon were seeded in 24-well plates and incubated with the different compounds for 24, 48 and 72 h. They were lysed in ice cold lysis buffer (Tris HCl 50mM, NaCl 100 mM EDTA 2 mM Triton-100 1% SDS 0.1%) containing protease inhibitors for 20 min. Cell lysates were collected and insoluble debris were removed by centrifugation. 20 μg of proteins were analyzed by western blotting using anti-NS5A and anti-β tubulin antibodies. Peroxidase-conjugated goat anti-mouse secondary antibody (Jackson Immunoresearch) was used for the revelation using ECL western blotting substrate (Thermo Fischer Scientific).

### Cell-to-cell transmission assay

Huh-7 cells were infected with HCVcc at MOI of 0.01. After 2 hours of contact, the inoculum was removed and replaced with fresh DMEM containing TF1, TF2, TF3 at 25 μg/ml or DMSO along with neutralizing antibody mAb 3/11 to prevent infection via the cell culture medium. The cells were propagated for another 72 h prior to immunofluorescence staining as described above. Quantification was done in 3 independent wells by taking independent pictures of different field of each well.

### Statistical Analysis

The statistical test used is a Kruskal Wallis nonparametric test followed by a Dunn’s multicomparison post hoc test with a confidence interval of 95% to identify individual difference between treatments. P values < 0.05 were considered as significantly different from the control. The data were analyzed using Graph Pad Prism (Version 5.0b) by comparisons between each treated group and untreated group (DMSO control).

## Results

### Theaflavins inhibit HCV infection

We and others have shown that EGCG, the major polyphenol present in green tea extract, was active against HCV infection (15,16,28), but no data are available on black tea polyphenols. In this context, three theaflavin derivatives, including TF1, TF2, and TF3, were extracted from CTC Assam black tea and identified by HPLC (Fig 1) with retention times of 10.49, 43.04, and 26.54 min, respectively. The average molecular weight of TF1 (564.49 g/mole), TF2 (716.59 g/mole) and TF3 (868.709 g/mole) was used for calculating the molar concentration of theaflavins which were generally used in micrograms in the experiments.

**Fig 1.**
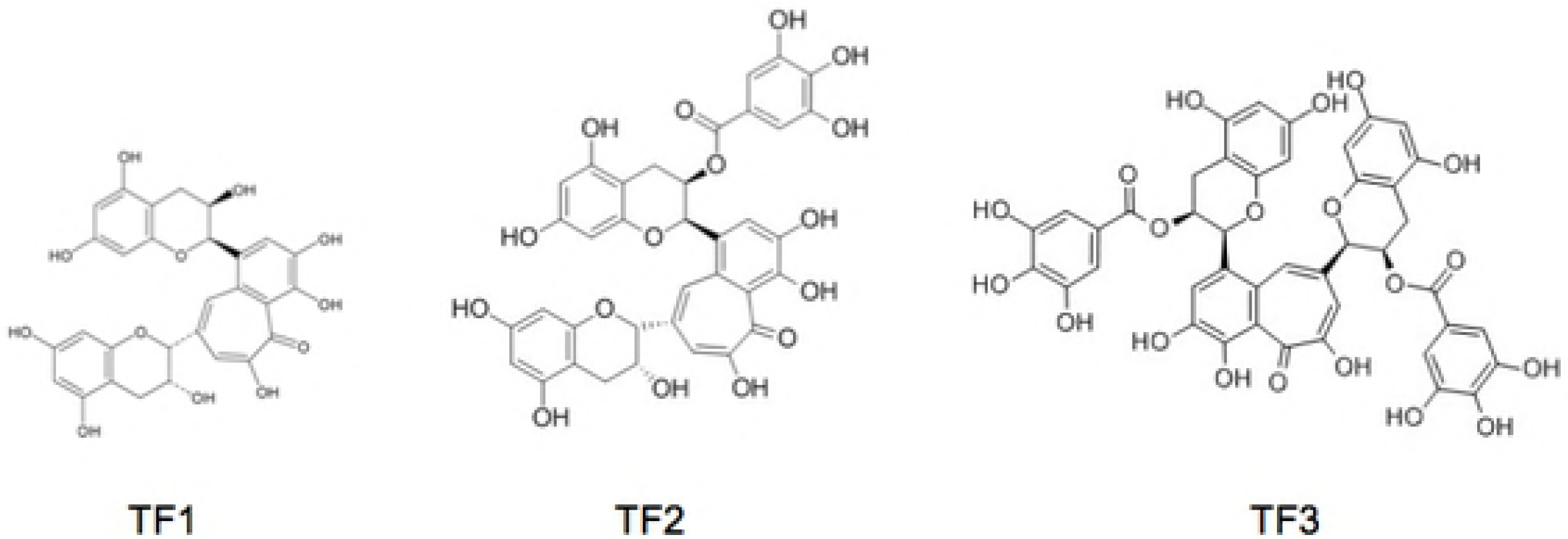
Chemical structures of theaflavins extracted from black tea. The inhibitory activity of theaflavins against HCV infectivity was carried out in dose-response experiment. Huh-7 cells were infected with HCV strain JFH1 in the presence of increasing concentrations of theaflavins, 0 to 100 μg/ml. Quantification of infected cells by immunofluorescence labeling of E1 envelope protein showed significantly decreased infectivity of HCV in the presence of each three theaflavins, with a clear dose-dependent inhibitory effect (Fig 2A). The half effective concentration (EC_50_) was calculated for each molecule to be 17.89, 4.08, and 2.02 μM for TF1, TF2 and TF3, respectively. TF3 is the most active. We then tested the effect of theaflavins on cell viability by treating Huh-7 with a range of theaflavin concentrations that were used in inhibition experiments, 0 to 250 μg/ml, with three different exposure times, 24, 48 and 72 h (Fig 2B-D). Theaflavins did not decrease cell viability up to 100 μg/ml after short (24 h) to long (72 h) exposure times, a concentration much higher than the active antiviral concentrations.

**Fig 2.**
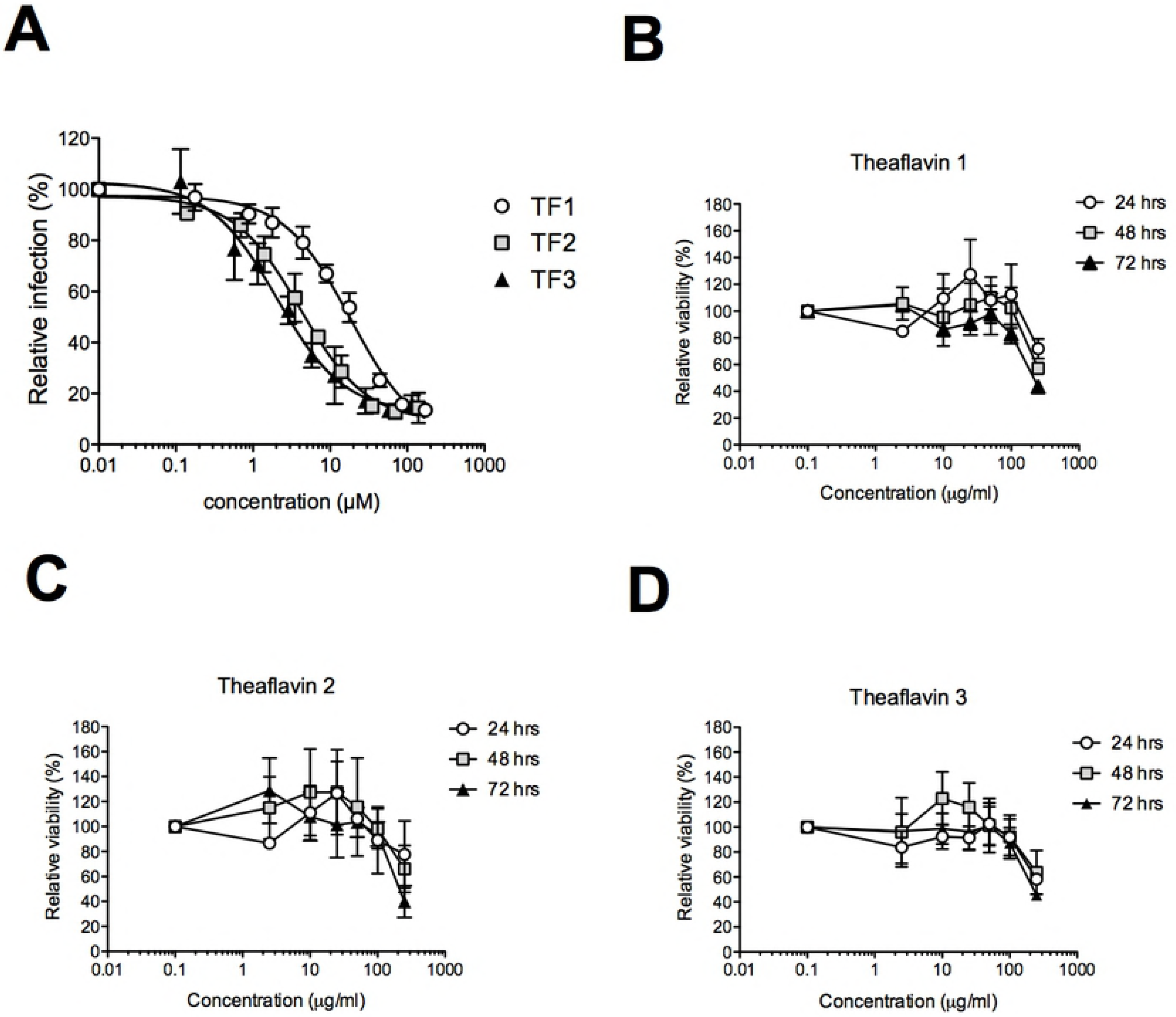
Theaflavins inhibit HCV infection. **(A)** Huh-7 cells were inoculated with HCV JFH-1 in the presence of given concentration of the different theaflavins. Inoculum was removed and replaced with medium containing theaflavins. Cells were fixed at 30 h post-infection and subjected to immunofluorescence to quantify number of infected cells. Results are expressed relative to DMSO control without theaflavins (100%). Huh-7 cells were incubated with given concentrations of TF1 **(B)**, TF2 **(C)** or TF3 **(D)** at different concentrations during 24, 48 or 72 h. Cell viability was measured using an MTS-based assay. Optical density at 490 nm was measured at the different time points. Results are expressed as mean ± SEM (error bars) of three independent experiments performed in triplicates and relative to control conditions without compounds.

### Theaflavins inhibit HCV entry and not replication

The HCV infection cycle is a multistep process that involves entry of the virus into the cell, replication of the genomic RNA and assembly/release of viral particles. To determine at which step the different theaflavins exert their action, a time of addition experiment was performed. The compounds were added before, during or after inoculation of HCV to Huh-7 cells. EGCG, an inhibitor of HCV entry, and Boceprevir, a HCV NS3 protease inhibitor, were used as controls. The result clearly demonstrates a significant inhibition of HCV only in the presence of theaflavins added during inoculation, like EGCG (Fig 3A). There was no inhibition when compounds were added to cells prior to the infection or post-infection. As expected, Boceprevir inhibits the post-inoculation step, corresponding to replication. These results, taken together, suggest that theaflavins act at an early step of the viral infectious cycle, most probably the entry step.

**Fig 3.**
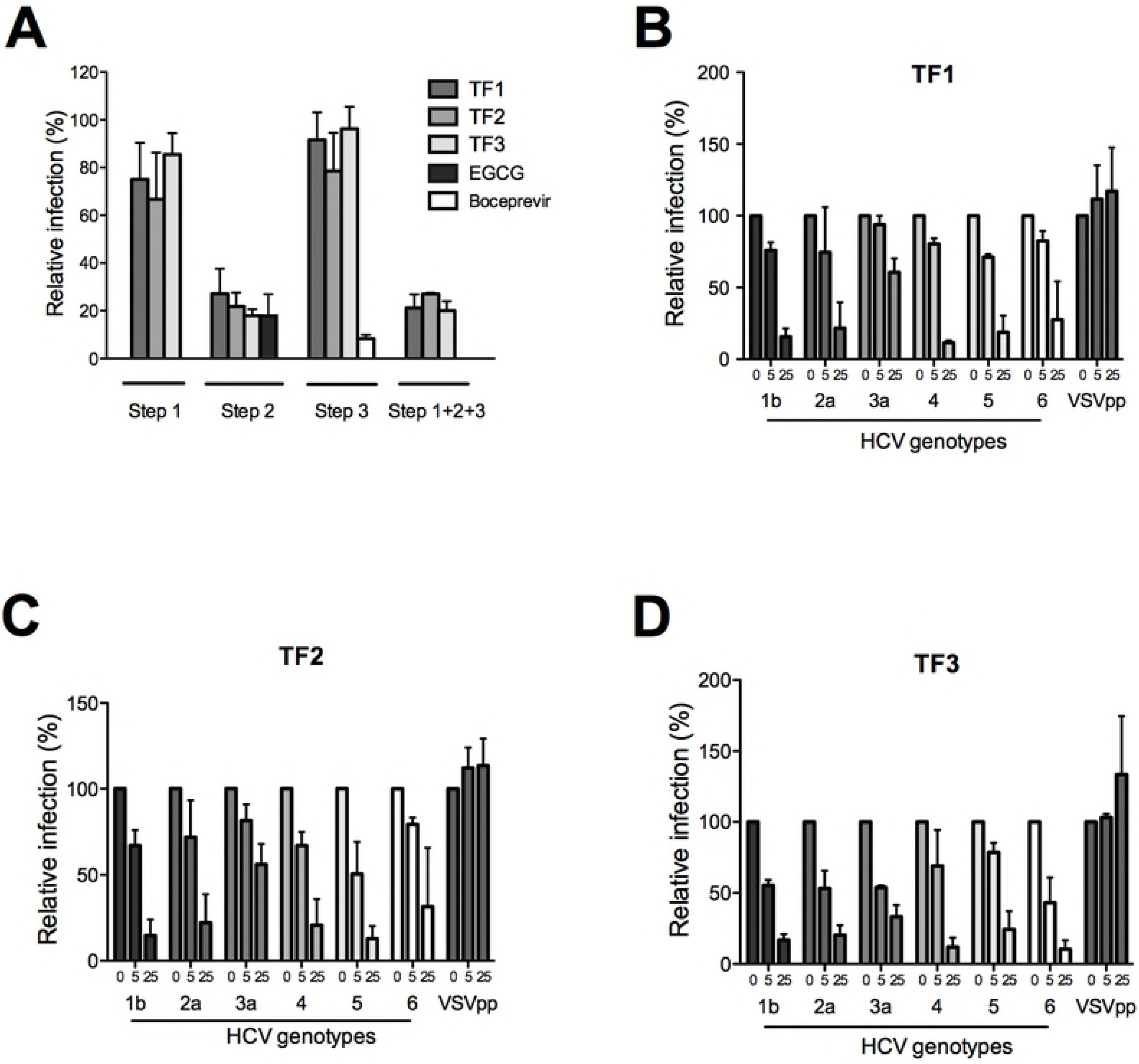
**Theaflavins inhibit HCV entry in a pan-genotypic manner.** (**A**) Huh-7 cells were inoculated with HCV JFH-1 for 2 h and fixed at 30 h post-infection to quantify the number of infected cells. Theaflavins, at 25 μg/ml were either added for 2 h to the cells before infection (step 1), or during inoculation (step 2), or during the 28 h after inoculation (step 3), or during the 3 steps (step 1+2+3). EGCG at 50 μM or Boceprevir at 1 μM were added either during the inoculation step or post-inoculation respectively. (**B-D**) Huh-7 cells were inoculated with HCVpp of the given genotypes or VSVpp in the absence (0) or presence of theaflavins at 2.5 or 25 μg/ml. Cells were lysed 48h after infection and luciferase activity quantified. Results are expressed as mean ± SEM (error bars) of 3 independent experiments performed in triplicate. Data are normalized to the DMSO, which is expressed as 100% infection.

Viral pseudoparticles are good models for entry inhibitor assay and also for screening against different viral genotypes (29). To confirm that theaflavins act at the entry step, HCVpp were produced and inoculated to Huh-7 cells in the presence of each theaflavins at active concentrations (EC_50_ and 10 × EC_50_). Envelope glycoproteins of different HCV genotypes were used, genotype 1b, 2a, 3a, 4, 5 and 6. The result shows that all the theaflavins inhibit infection of HCVpp in a dose dependent manner (Fig 3B-D) confirming their activity on HCV entry. Moreover, the antiviral activity of each theaflavin is pan-genotypic, with an inhibition of infection for all HCV genotype tested. HCVpp of genotype 3a are less inhibited by theaflavins than HCVpp of other genotypes, even if TF3 seems more active on this genotype than TF1 or TF2.

Results presented in Fig 3A suggest that theaflavins are not active on HCV replication because no inhibitory effect was observed when the compounds were added during the 28 h postinoculation. To confirm this data, HCV subgenomic replicon containing a GFP-tagged NS5A was used. Replicon cells were incubated with TF1, TF2 or TF3 at 25 μg/ml, or Boceprevir at 1 μM, for 72 h. NS5A protein quantified by Western blot analysis. The result shows that the replication of the replicon is not affected by the presence of any of the theaflavins but is inhibited by Boceprevir (Fig 4). Taken together, our results show the TF1, TF2 and TF3 are inhibitors of HCV entry and not replication.

**Fig 4.**
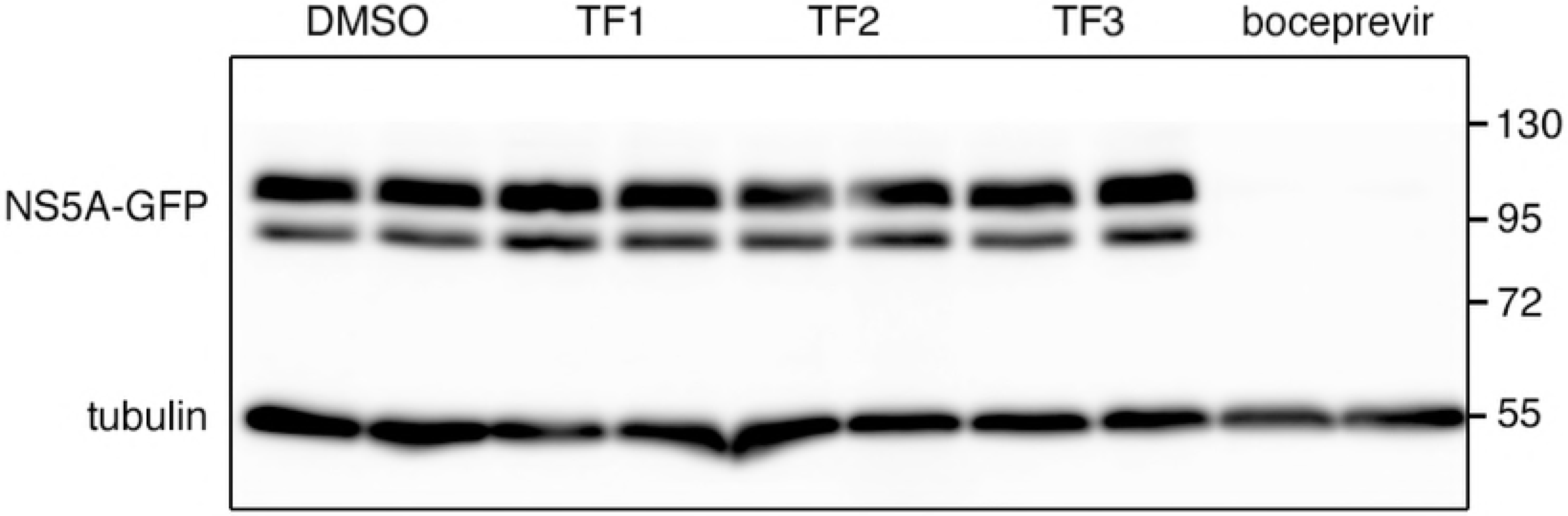
Theaflavins are not inhibitors of HCV replication. Huh-7 cells expressing a HCV replicon containing a GFP-tagged NS5A were incubated in the absence (DMSO) or presence of theaflavins at 25 μg/ml, or Bocebrevir at 1 μM, for 72 h. Cells were lysed and Western blot analysis was performed to detected NS5A-GFP and tubulin with specific antibodies. Data are representative of 3 independent experiments.

### Theaflavins act directly on the viral particle

Theaflavins can inhibit HCV entry by acting either directly on the viral particle or on cellular factors. This second hypothesis seems unlikely because no effect of theaflavins on HCV infection was observed when cells were pre-treated with the molecules before infection (Fig 3A). To determine if theaflavins act directly on the viral particle, HCV viral stocks were incubated with the compounds at 25 μg/ml prior inoculation. The viral stocks containing the compounds were then diluted 10 times before inoculation leading to a final concentration of theaflavins of 2.5 μg/ml for the inoculation step. In parallel, cells were inoculated with a non-treated virus in the presence of the compounds at the two different concentrations, 2.5 and 25 μg/ml. As shown in Fig 5, the inhibitory effect of theaflavins on pre-incubated virus is similar to the one observed when virus is inoculated on cells at high concentration (25 μg/ml), demonstrating that theaflavins act directly on HCV particle before entry.

**Fig 5.**
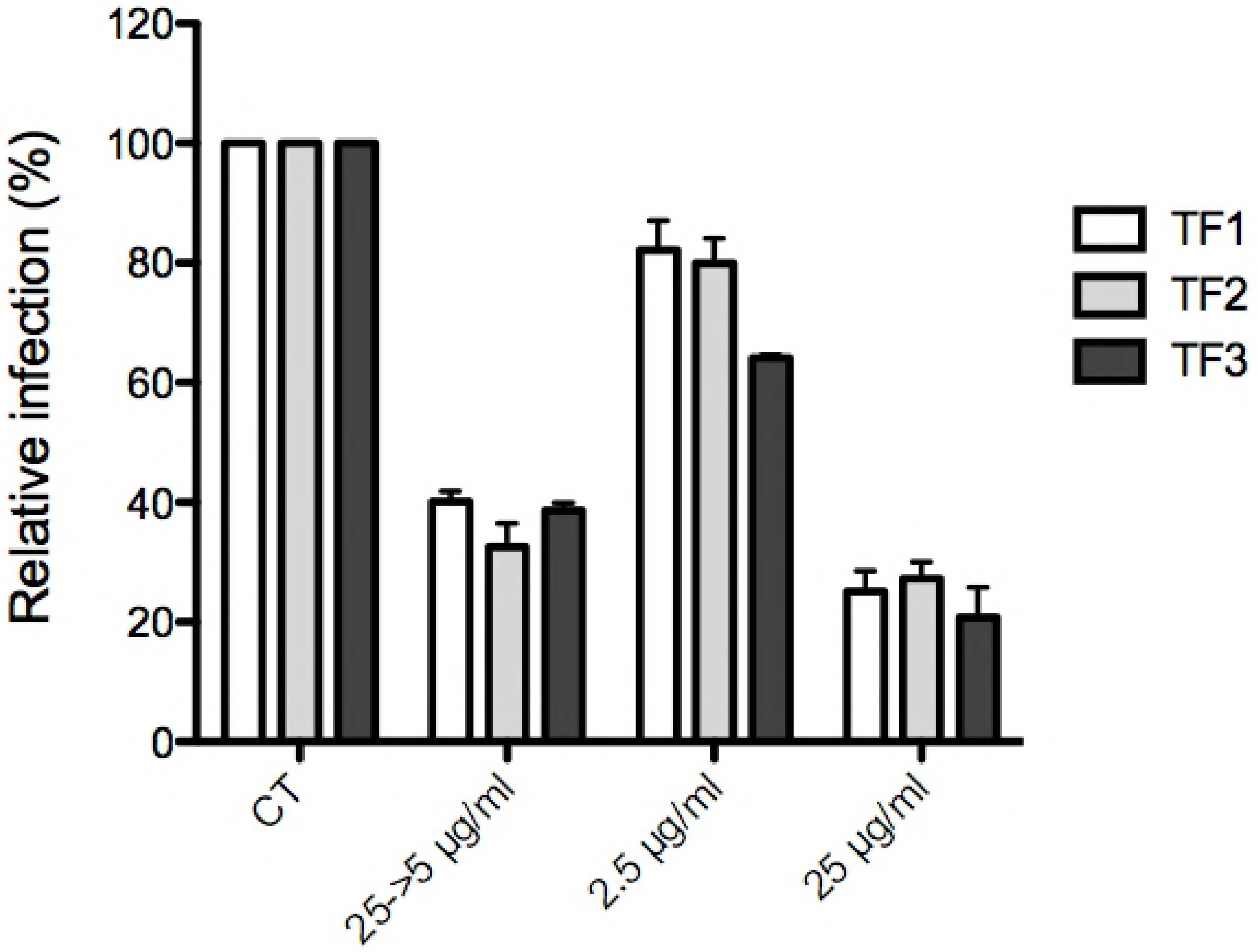
Theaflavins act on the viral particles prior inoculation. HCV virus stock was pre-incubated with 25 μg/ml of theaflavins for 1 h before inoculation, and then the mixture was diluted 10 times before being used for the inoculation, leading to a final concentration of theaflavins of 2.5 μg/ml. In parallel, Huh-7 cells were inoculated with HCV in the presence of theaflavins either at 2.5 or 25 μg/ml. Cells were fixed and subjected to immunofluorescence to quantify infected cells as described above. Data are normalized to the DMSO, which is expressed as 100% infection. The results are presented as means ± SEM of three independent experiments performed in triplicate.

### Theaflavins inhibit HCV cell-to-cell propagation

HCV infection can follow two different modes of transmission, either by direct entry into the cell via extracellular medium or via cell-to-cell spread between two neighboring cells. This last mode of transmission is resistant to neutralizing antibodies and seems to be a major way of HCV infection (30). Therefore, it is important to identify antiviral agents able to block cell-to-cell transmission. To determine if theaflavins can inhibit HCV cell-to-cell spread, Huh-7 cells were infected with HCVcc with no inhibitor, at low MOI to visualize foci, and further incubated with theaflavins in the presence of neutralizing antibodies to prevent extracellular infection. 72 h postinfection, the number of cells in the foci were quantified. The results shows that TF1, TF2 and TF3 significantly reduce number of cells in the foci meaning reduction in cell-to-cell spread (Fig 6), like EGCG a known inhibitor of cell-to-cell transmission (15). This shows that theaflavins are potent inhibitors of both HCV modes of entry.

**Fig 6.**
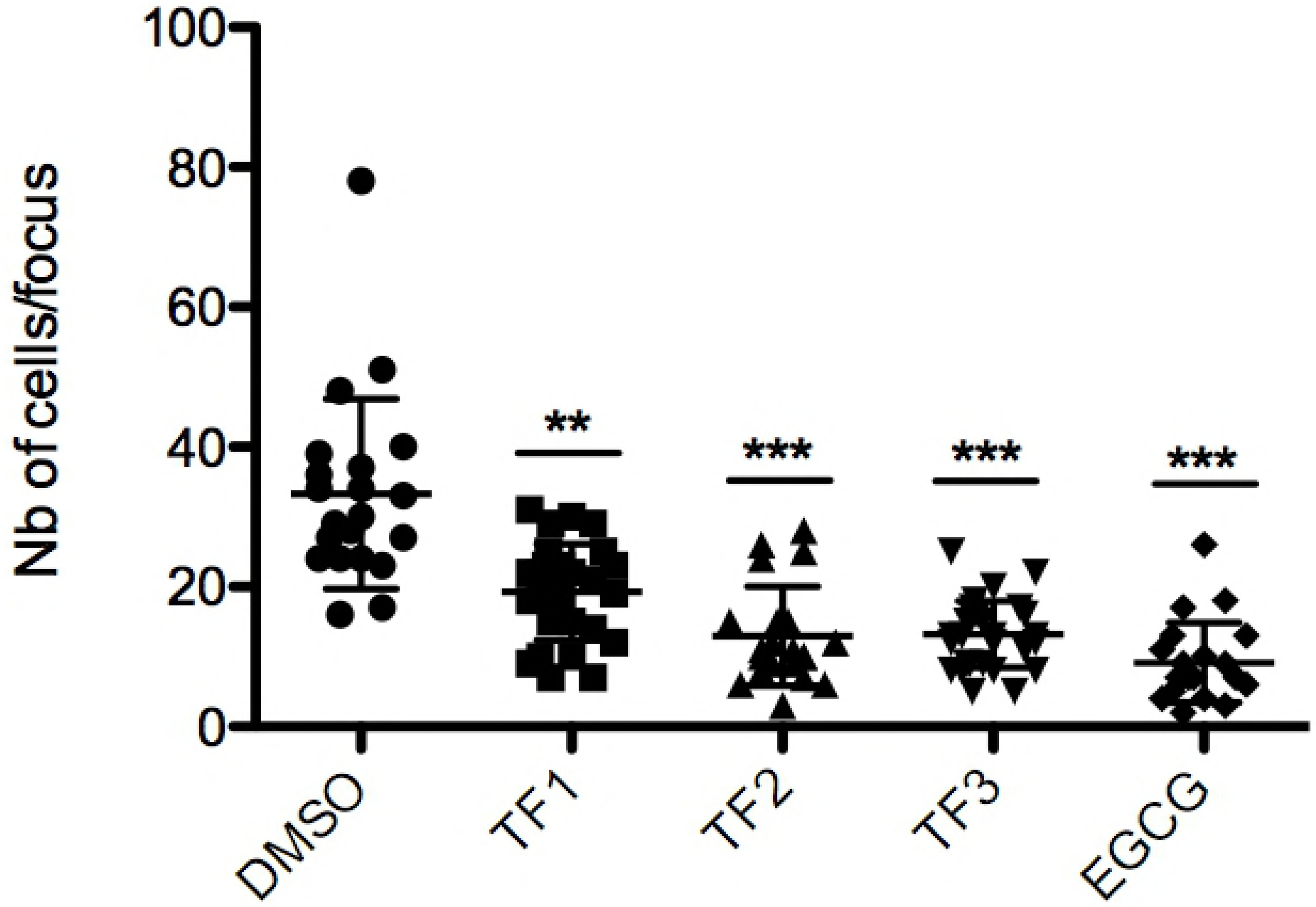
Theaflavins inhibit cell-to-cell spread. Huh-7 cells were inoculated with HCV JFH-1 for 2 h. Inoculum was replaced by medium containing mAb 3/11 neutralizing antibody to prevent extracellular propagation. Theaflavins at 25 μg/ml or EGCG at 50μm were added in the medium after inoculation. Cells were fixed 72 h after infection and subjected to immunofluorescence detection of E1 HCV envelope protein as described. The number of cells/focus of 3 independent fields of three independent wells was quantified. Error bars represent SD. Data are representative of 3 independent experiments. Statistical analysis were performed with the Kruskal Wallis nonparametric test followed by a Dunn’s multicomparison post hoc test. **, P < 0.01; ***, P < 0.005.

### Combination of theaflavins with replication inhibitors used in hepatitis C therapy

Finally, in order to determine the potential use of these compounds in antiviral therapy, we interrogated the effect of theaflavins in combination with the known DAA Sofosbuvir and Daclatasvir, two inhibitors of HCV replication that target HCV proteins NS5B and NS5A respectively (31,32). In the experiment, TF3, the most active theaflavin, was added at different concentrations along with Sofosbuvir or Daclatasvir at fixed concentration during infection of Huh-7 cells with HCVcc. The result shows that TF3 can increase the antiviral activity of both Sofosbuvir and Daclatasvir in an additive manner (Fig 7) demonstrating that it could be use in combination with DAA used in hepatitis C therapy.

**Fig 7.**
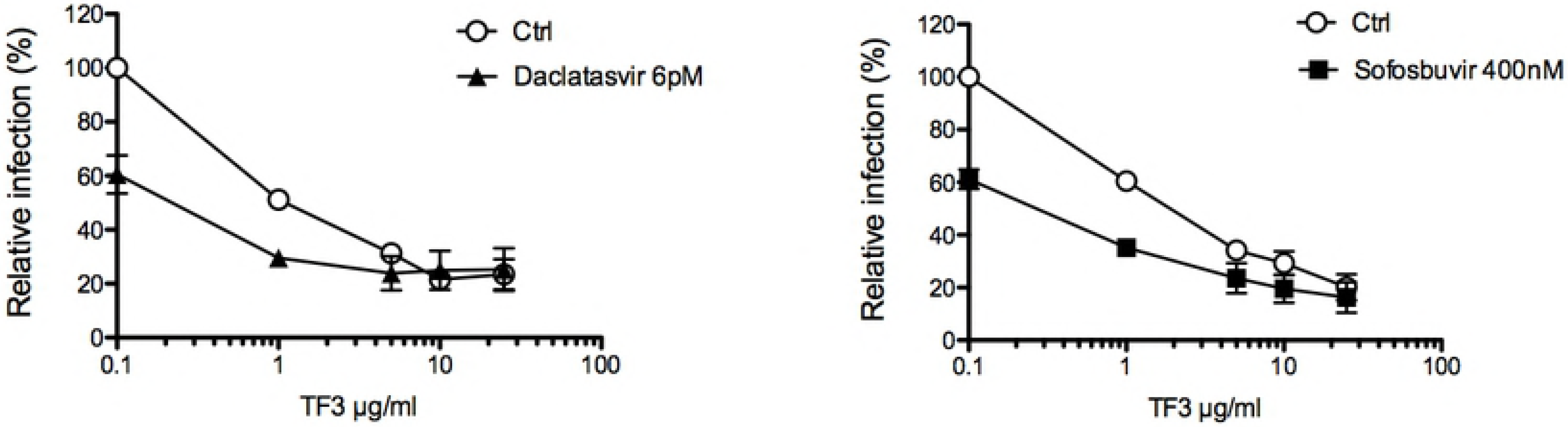
Theaflavins can be used in combination with DAA. Huh-7 cells were inoculated with HCV JFH-1 in the presence of TF3 at given concentrations. Inoculum was removed and replaced by medium containing or not either Daclatasvir at 6 pM or Sofosbuvir at 400 nM. Cells were fixed at 30 h post infection and subjected to immunofluorescent detection of E1 envelope protein. Results are expressed as mean +/− SEM (error bars) of 3 independent experiments performed in triplicate. Data are normalized to DMSO, which is expressed as 100% infection.

## Discussion

The present study identified new entry inhibitors of HCV, theaflavins, the major black tea polyphenols. Theaflavins inhibit all HCV genotypes and most probably induce inactivation of the viral particle before inoculation. Interestingly, they are also able to inhibit cell-to-cell spread of HCV, a major route of infection allowing evasion of the virus from neutralizing antibodies (30). Moreover, an additive action of TF3 in combination with molecules used for hepatitis C therapy was observed.

HCV is transmitted between hepatocytes via classical cell entry but also uses direct cell-to-cell transfer to infect neighboring hepatocytes. Cell-to-cell transmission was also shown to play an important role in dissemination and maintenance of resistant variants in cell culture models (33). HCV entry is highly orchestrated and essential in initiating viral infection and spread. This step represents a potential target for HCV inhibitors (34,35) but, up to now, no HCV entry inhibitor has been marketed, and few are in preclinical trial. The currently used therapy of DAA regimen cures more than 90% of infected patients, but the appearance of viral resistance, and recurrence of infection particularly in transplant patients is still a major limitation (36). Entry inhibitors given along with the DAAs would be expected to exhibit a synergistic effect (34,35). The present study supported this hypothesis when theaflavin and Sofosbuvir or Daclatasvir were added in combination and showed additive effect against HCV. This may lead to pivotal implication in therapeutic regimen of HCV especially because Sofosbuvir, in combination with other drugs, is a part of all first-line treatments for HCV, and also of some second-line treatments (36).

HCV treatment efficacy is influenced by infected viral genotype; therefore, treatment regimen is dependent on genotypes infected. Our study with HCVpp of different genotypes (1–6) showed dose-dependent inhibition of HCV by theaflavins (Fig 3B-D). This represents an important hit for further drug development.

In recent times, a number of promising natural products with anti-HCV activities have been discovered (37–39). They are of different origins and chemical structures and exert their antiviral effects at different levels within the virus life cycle. While green tea polyphenols have received the most attention in the past years, data presented in this study suggest that black tea theaflavins may also be potent candidates for antiviral drug development particularly as entry inhibitors against HCV. Our data show that TF3 has the most prominent effect against HCV in comparison to TF1 and TF2. TF3 (EC_50_ = 2.2 μM) also is more potent than EGCG (EC_50_ = 10.6 μM) (14).

Our results clearly indicate that all the three derivatives of theaflavin act directly on viral particles and may prevent cell surface attachment or receptor binding. In our earlier study cryo-electron microscopy imaging showed that EGCG and delphinidin, an anthocyanidin, have a bulging effect on HCV envelope of HCVpp (14). We might speculate that theaflavins might have similar effect as both polyphenols shared close structural similarity. However, the exact mechanism of action of theaflavins on HCV infection needs further studies.

TF3 seems to be most promising candidate against HCV and its efficacy may be attributed to the presence of double galloyl moiety. The same has been found for HSV-1 where TF3 was reported to be most effective (18). In contrast, the presence of the galloyl group in TF3 is not necessary for antiviral properties against calcivirus where all three theaflavin derivatives have similar efficacy (17). To understand the role of functional groups and galloyl moiety, further study is required through the approach of combinatorial and synthetic chemistry. It is interesting to note that theaflavins and EGCG have been described to inactivate the same viruses, HSV-1, HIV-1 and influenza virus (18–20,40). Taken together with the results presented here, it seems that these polyphenols from tea, black or green, have a very specific mode of action on enveloped viruses that could be more exploited in the context of antiviral therapy.

A future prospect of our study may be to determine the bioavailability of theaflavins in human liver. Henning et al performed study on liver tissue of mice after black tea consumption and showed that theaflavin relative absorption in liver is twice higher than EGCG (41). TF1 concentration in mice liver is estimated at 0.4 nmol g^−1^ tissue, approximately 0.4 μM. In contrast, TF2 and TF3 are poorly detected. Theaflavin content may vary from one tea to another. It has been shown that Assam black tea contains more theaflavins than Darjeeling tea, and particularly TF3 (125.9 and 26.8 μg/ml respectively) (42), which could lead to higher relative absorption. Even if the tissue concentrations of theaflavins are lower than their active concentrations, their use in combination with other DAA might reduce their EC_50_ as shown in Fig 7. Taken together, these results may be of crucial importance in developing a prophylactic drug for HCV risk group population.

In conclusion, the present study identified theaflavins as new entry inhibitors of HCV infection. Their pan-genotypic action and ability to inhibit cell-to-cell spread are major advantages for further evaluation for drug development, as well as their efficacy in combination with actual antiviral therapy. Moreover, their ease of extracting, availability and popularity of tea as a drink make them interesting candidate as entry inhibitors against HCV infection.

## Acknowledgements

The authors are thankful to Department of Health Research, Ministry of Health and Family welfare, Government of India, for providing post-doctoral overseas training fellowship to P. Chowdhury. The authors are also thankful to Director, TTRI and Secretary, TRA for their support to implement the research fellowship grant.

